# Geometry-based navigation in the dark: Layout symmetry facilitates spatial learning in the house cricket, *Acheta domesticus*, in the absence of visual cues

**DOI:** 10.1101/2019.12.28.886655

**Authors:** Bartosz Baran, Michał Krzyżowski, Zoltán Rádai, Jacek Francikowski, Mateusz Hohol

## Abstract

The capacity to navigate by layout geometry has been widely recognized as a robust navigational strategy. It was reported in various species, albeit most studies were performed with vision-based paradigms. In the presented study, we aimed to investigate layout symmetry-based navigation in the house cricket, *Acheta domesticus*, in the absence of visual cues. For this purpose, we used a non-visual paradigm modeled on the Tennessee Williams setup. We also verified the inaccessibility of visual cues for tested insects using antennal positioning reflex towards looming stimulus and by testing the performance of blinded crickets. In the main experiment, we tested the crickets’ capacity to learn to find a cool spot positioned centrally in heated arenas of different shapes (i.e., circular, square, triangular, and asymmetric quadrilateral). We found that the symmetry of the arena significantly facilitates crickets’ learning to find the cool spot, indicated by the increase of time spent on the cool spot and decrease of the latency of locating it in subsequent trials. To investigate possible mechanisms utilized by crickets during the experiment, we analyzed insects’ approach paths to the spot. The trajectories were grouped in four distinct clusters corresponding to both heuristic and directed strategies of approaching the target, with the dominance of a semi-directed strategy (thigmotactic phase preceding direct navigation to the target). Against these results, we discuss the possibility of insects’ navigation by using a non-visual space representation and possible limitations of navigation capacities in such conditions in relation to multimodally-guided navigation.

## 1. Introduction

Spatial navigation plays a vital role in the lives of the animals, allowing them to successfully forage for food, find their way back to nests, or localize mating spots. To this end, animals employ a spectrum of strategies that allow them to repeatedly return to the memorized locations despite the constantly changing environment and disrupting stimuli (Gallistel 1990; Thinus-Blanc et al. 2010; Tommasi et al. 2012). One of the strategies enabling the mitigation of the constant change of environmental features is to navigate by the relational pattern of the objects in the surrounding space (layout geometry) instead of by their particular features. Those relations could be perceived via one or more modalities (Cheung et al. 2008; Stürzl et al. 2008; Webb and Wystrach 2016). Nevertheless, the contribution of vision seems to be investigated much more than any other modality, constituting a somewhat “visiocentric” bias in spatial navigation studies (Hohol et al. 2017). However, the factual capacity for processing geometric relations should be feature-independent, as these relations are preserved across the modalities (Duval 2019). In this regard, we want to highlight the importance of conducting studies focused on non-visual modalities. This approach follows similar rationality as the cross-modal testing of abstract numerical representations (Izard et al. 2009; Dehaene 2011; Butterworth 2022).

The body of research on animals’ navigation by layout symmetry consists primarily of data obtained from vertebrates. It was found that using layout geometry as a cue for navigation is not task-dependent since vertebrates are able to localize the center of an arena based on its overall geometric shape and are capable of transferring this knowledge to other geometrically regular enclosures (Tommasi et al. 1997; Gray et al. 2004; Tommasi and Thinus-Blanc 2004; Tommasi and Giuliano 2014). Localizing the center based on discrete landmarks was also investigated (Kamil and Jones 1997, 2000; Sutton et al. 2000; Potì et al. 2010). Research on insect navigation utilizing environmental geometry is scarce and mainly concerns navigation in rectangular arenas (Wystrach and Beugnon 2009; Sovrano et al. 2012; Lee and Vallortigara 2015). However, aside from the studies directly concerning layout geometry, there are other reports suggesting that miniature nervous systems are able to process geometric properties of objects, such as symmetry (Giurfa et al. 1996; Møller and Sorci 1998; Rodríguez et al. 2004; White and Kemp 2020).

While there is still no consensus about the exact mechanism responsible for the observed behavioral patterns of layout geometry-driven navigation in vertebrates (Cheng et al. 2013; Sutton and Newcombe 2014; Duval 2019; Hohol 2020), in the case of insects, it is widely accepted that view matching (VM) is the core mechanism behind this mode of spatial navigation (Wehner et al. 1996; Collett and Rees 1997; Judd and Collett 1998; Wystrach et al. 2011; Wystrach and Graham 2012b, a; Collett et al. 2013). In brief, the VM approach assumes that the animal records a visual snapshot of the area surrounding the goal and then moves so as to minimize the discrepancy between the snapshot and the actual view. Wherein the memorized “view” is not simply understood as a mental image but instead as a set of encoded parameters including depth, motion, edges, or specific features.

Geometry-driven navigation may be constituted by different perceptual and cognitive mechanisms, depending on the species (Vallortigara 2018). For instance, *Gigantiops destructor*, a neotropical formicine ant tested by Wystrach and Beugnon (2009) for the presence of rotational errors, is an animal highly dependent on vision. Therefore, the VM-based explanation of its navigational behavior in geometrically regular arenas is convincing. Nevertheless, this does not automatically imply that the navigation of other insects, such as adult house crickets, which are predominantly nocturnal animals (Cymborowski 1973; Górska-Andrzejak and Wojtusiak 2003), in the aforementioned enclosures would also be driven by VM. Although geometry is considered mainly as a vision-based phenomenon, it has been demonstrated that geometric cognition in various species, both regarding objects (Marlair et al. 2021), and layouts (Sovrano et al. 2018, 2020; Nardi et al. 2021), could be grounded also in other modalities, e.g., proprioceptive and kinesthetic (Alary et al. 2008; Sguanci et al. 2010; Marlair et al. 2021). Aside from navigation by visual cues (Doria et al. 2019), crickets have been tested in experiments involving auditory information (Reeve and Webb 2003; Poulet and Hedwig 2005). Nevertheless, none of the existing studies allow us to infer their ability to use non-visually perceived layout geometry for spatial learning and navigation.

In the present study, we tested whether house crickets (*Acheta domesticus*) are able to find a target positioned centrally in an arena in the absence of visual cues, and if so, whether the symmetry of the spatial layout facilitates learning of such a task. To this end, we employed a variant of the center-finding paradigm, where the animal is required to find an inconspicuous cool spot positioned at the center of the following experimental enclosures: circular, square, equilateral triangular, and asymmetric quadrilateral. Originally, the task was developed by Tommasi, Vallortigara, and Zanforlin (1997) to test the spatial cognition of chickens, and later it was used to investigate other vertebrate species, namely, pigeons (Gray et al. 2004), rats (Tommasi and Thinus-Blanc 2004), and human children (Tommasi and Giuliano 2014).

Aside from exploring the possible vision-independent geometric navigation, the non-visual testing conditions meet the ecological validity standard as the house cricket imagoes are predominantly active at night (Cymborowski 1973; Górska-Andrzejak and Wojtusiak 2003). As the VM generally explains navigational behavior in a low-level way, namely, the overall encoding of the layout geometry is not required, we prevented the insects from using view-based place finding. For this purpose, we employed a non-visual paradigm modeled on the Tennessee Williams (TW) setup (Mizunami et al. 1998; Wessnitzer et al. 2008), which is a “dry” analog of the Morris (1981) water maze (MWM) test commonly used for assessing navigational capabilities. The unavailability of visual cues was assessed prior to the main experiment by testing the occurrence of antennal positioning reflex towards the looming stimulus under the illumination used for the center finding assessment. Moreover, for further confirmation of the non-visually driven nature of observed effects, the center-finding in crickets with enamel-covered eyes was tested. Additionally, measures were taken to prevent the potential influence of olfactory and auditory stimuli, which could provide insects with cues for reorientation.

We expected to find that the behavior observed in this study would converge with those previously reported in vertebrates (Tommasi et al. 1997; Gray et al. 2004; Tommasi and Thinus-Blanc 2004; Tommasi and Giuliano 2014). First of all, we hypothesized that crickets would be able to learn to find the centrally located cool spot. Since geometrically regular layouts are easier to navigate, our second hypothesis was that cricket would learn to find the center more efficiently in conditions with symmetric arenas (circular, square, and triangular) compared to the asymmetric quadrilateral one.

## 2. Materials and methods

### 2.1. Experimental setup

In the presented study, the center-finding paradigm (Tommasi et al. 1997), combined with the non-visual variation of the TW setup (Wessnitzer et al. 2008). Note that Wessnitzer et al.’s setup was actually adapted from the study by Mizunami et al. (Mizunami et al. 1998), and more recently used by Ofstad, Zuker, Reiser (Ofstad et al. 2011).

The experimental apparatus (Fig. 1 and Fig. 2) consisted of a leveled, matted, white glass sheet and variously shaped arenas (circular, square, triangular and asymmetric quadrilateral), all of the same height (25cm) and adjusted to approximately the same area (709±3%cm^2^ - circular d=30, square: a=27, triangular: a=40 asymmetric quadrilateral: a=37 b=24 c=23 d=26 α=67° β=80° γ=100° δ=113° [cm]). The arenas were made of solvent-welded white Lucite and devoid of any visual cues. Solvent-welds were utilized to ensure that corners don’t provide any attachment point for the crickets. Thus, even if leaning against the corner occurred it was brief as the insect slid back inside the arena. The surface of the glass was uniformly heated to 50±1°C with IR heating lamps (4 × 250W bulbs, heat distribution was evaluated with the FLIR T640 thermal camera) with the exception of a centrally localized cool spot that was maintained at a constant temperature of 25±1°C, preferable by *A. domesticus* (Lachenicht et al. 2010), by a water-cooling block (ø60mm). Exterior parts of the arenas were covered with aluminum foil to ensure that the walls were at least partially heated. All experimental trials were performed in a soundproofed dark room. Arenas were illuminated with a red LED ring (24 × WS2812B) emitting red light with a wavelength of 620-625nm, which is below the detection threshold of the retinal receptors of crickets (Herzmann and Labhart 1989), and thus was chosen in order to ensure the lack of visual cues. Before each trial, the enclosures of the arenas were rotated by 45°, and the whole setup was thoroughly wiped with 70% (v/v) denatured ethyl alcohol to eliminate any olfactory clues.

**Figure 1.**
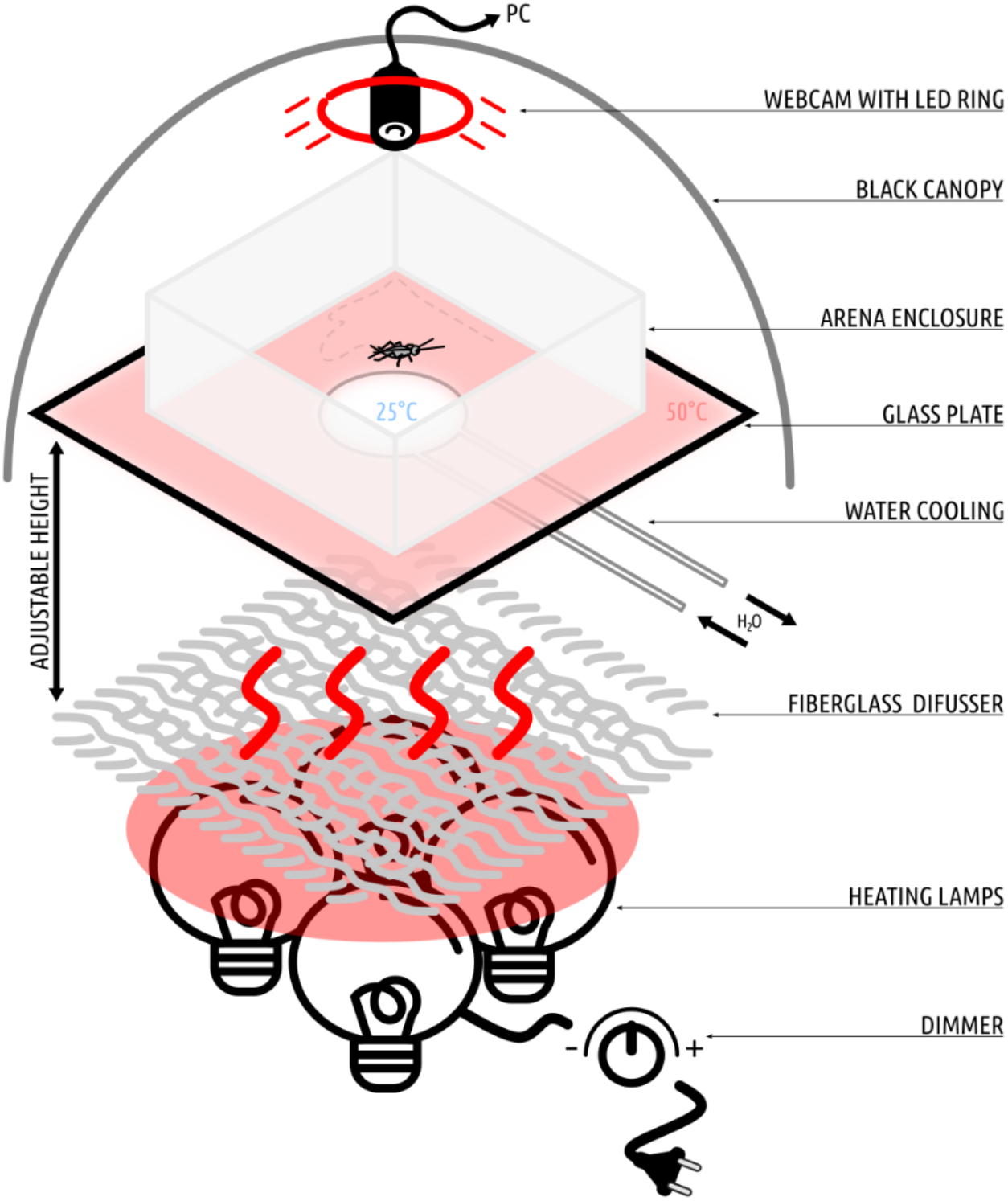
The applied experimental setup iterates the TW setup, namely a spatial learning task similar to the MWM, where the insect explores a plate heated to an aversive temperature to find the inconspicuous cool spot on which it can rest. Our apparatus consisted of 4 × 250W dimmable heating lamps with a fiberglass cloth diffuser mounted above them. Thermal radiation generated by the lamps uniformly heated the bottom side of the glass plate (500×500×4mm), painted with two layers of heat-resistant enamel. The first layer was made of white enamel (to provide a contrasting background for insect tracking), and the second was painted black (for thermal absorption). On the backside of the plate, the 3D printed water cooler (ø60mm) was attached with a Gecko pad and connected to a continuous flow of cool water adjusted to ensure a constant temperature of a cool spot on the surface of the plate. The glass was chosen due to its low thermal conductivity, which allows for the creation of a sharp thermal boundary around the cool spot. The upper surface of the glass was matted to ensure sufficient traction for the insects. Differently shaped arena enclosures were placed on the surface of the glass. The setup was calibrated with the aid of thermal imaging to provide stable temperatures of 50±1°C on the hot part and 25±1°C on the cool spot (see Fig. 2). Prior to every training session, the setup was warmed to the desired temperatures. For each arena shape, 15 crickets were tested (n = 15 × 4).

**Figure 2.**
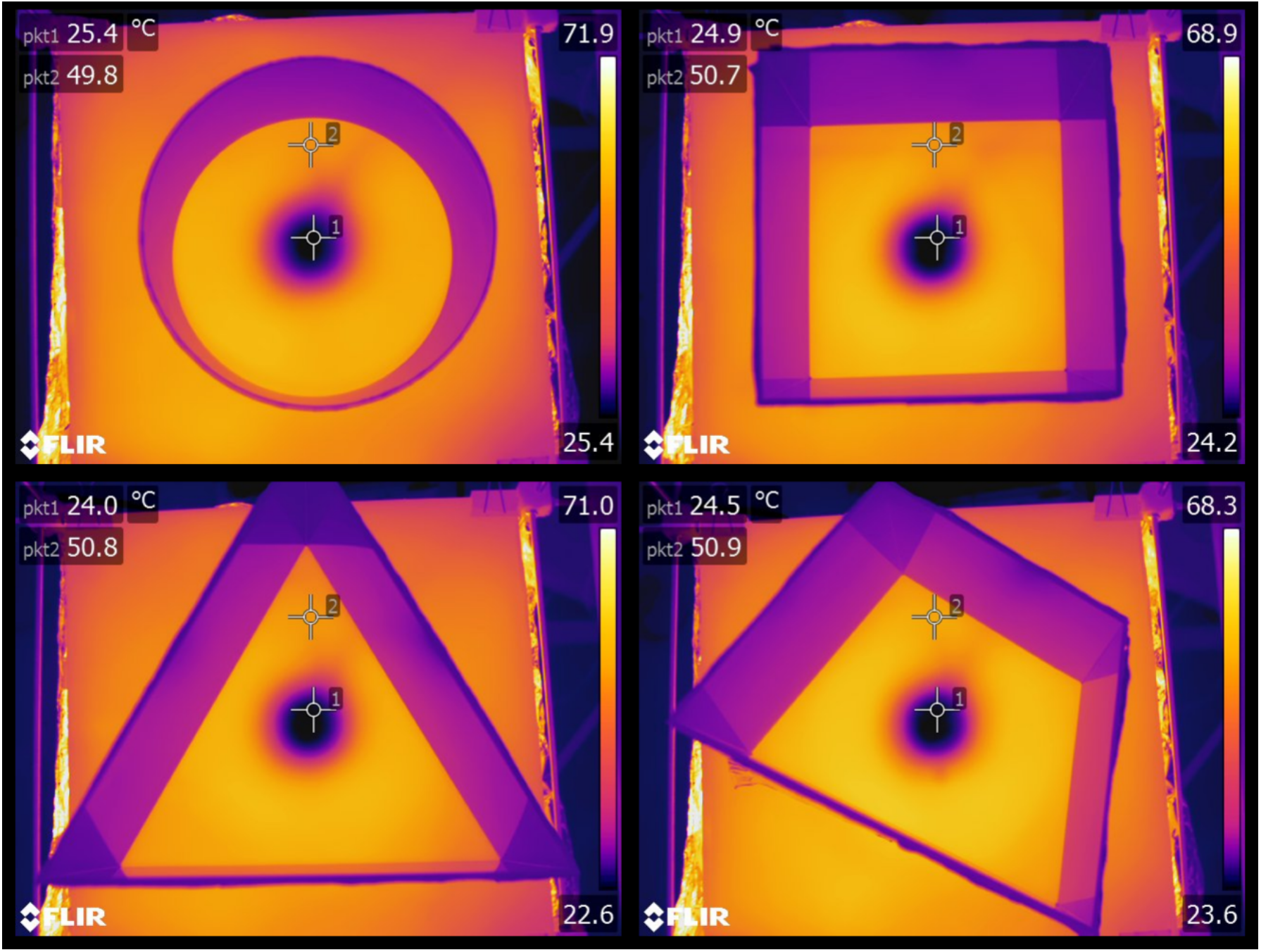
Thermal imaging of the heat distribution on the surfaces of the arenas. Two measurement points, pkt1 and pkt2, indicate the temperatures of the cool spot and the rest of the arena, respectively. Images were acquired with a FLIR Systems AB high-performance thermal camera FLIR T640 (640×480px IR resolution, sensitivity 0.04°C at 30°C).

**Figure 3.**
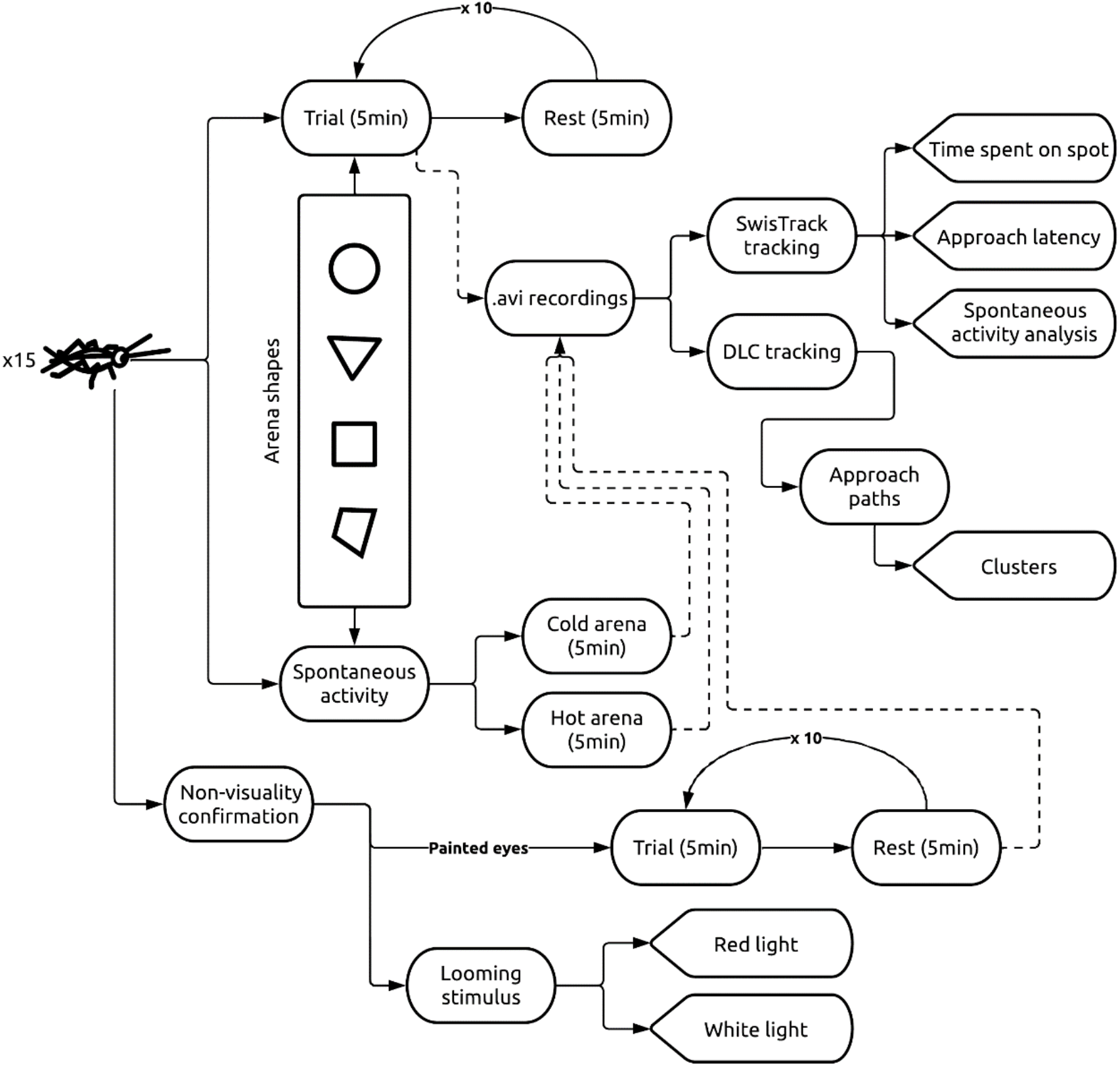
Overview of the entire study. The learning procedure was based on a study by Wessnitzer, Mangan, and Webb (2008) consisting of 10 × 5 min long trials alternating with 5 min rests. The duration of the rest time was adjusted from 2 min used by Wessnitzer et al. (2008) to reduce elevated erratic escape behavior, which was observed in a pilot study conducted prior to the primary experiment.

### 2.2. Animal husbandry

House crickets (*Acheta domesticus*, wild type) used in the study were acquired from a stock colony maintained at the Institute of Biology, Biotechnology and Environmental Protection of the University of Silesia in Katowice. Insects were reared under constant conditions of 30±2°C, 40±10% RH, and a 12:12 light:dark photoperiod with water and food pellets *ad libitum*. In all trials, healthy adult (1-2 days after the imaginal molt) male crickets were used. After conducting experimental assessment crickets were discarded to separate retirement colony, so that each individual participated only once in the experiments.

### 2.3. Confirmation of inaccessibility of visual cues

To confirm the inaccessibility of the visual cues in the main experiment, we employed a two-way approach. Firstly, we ensured that the used setup sufficiently suppressed the usage of the visual cues. To this end, we conducted a test of visual behavior *per se* (antennal positioning reflex) under the same illumination as was used during the main experiment (Fig. 4). Secondly, we conducted an additional test with blinded insects (eyes carefully painted over with opaque blue Edding 751 paint marker) using exactly the same procedure and conditions as in the main experiment.

**Figure 4.**
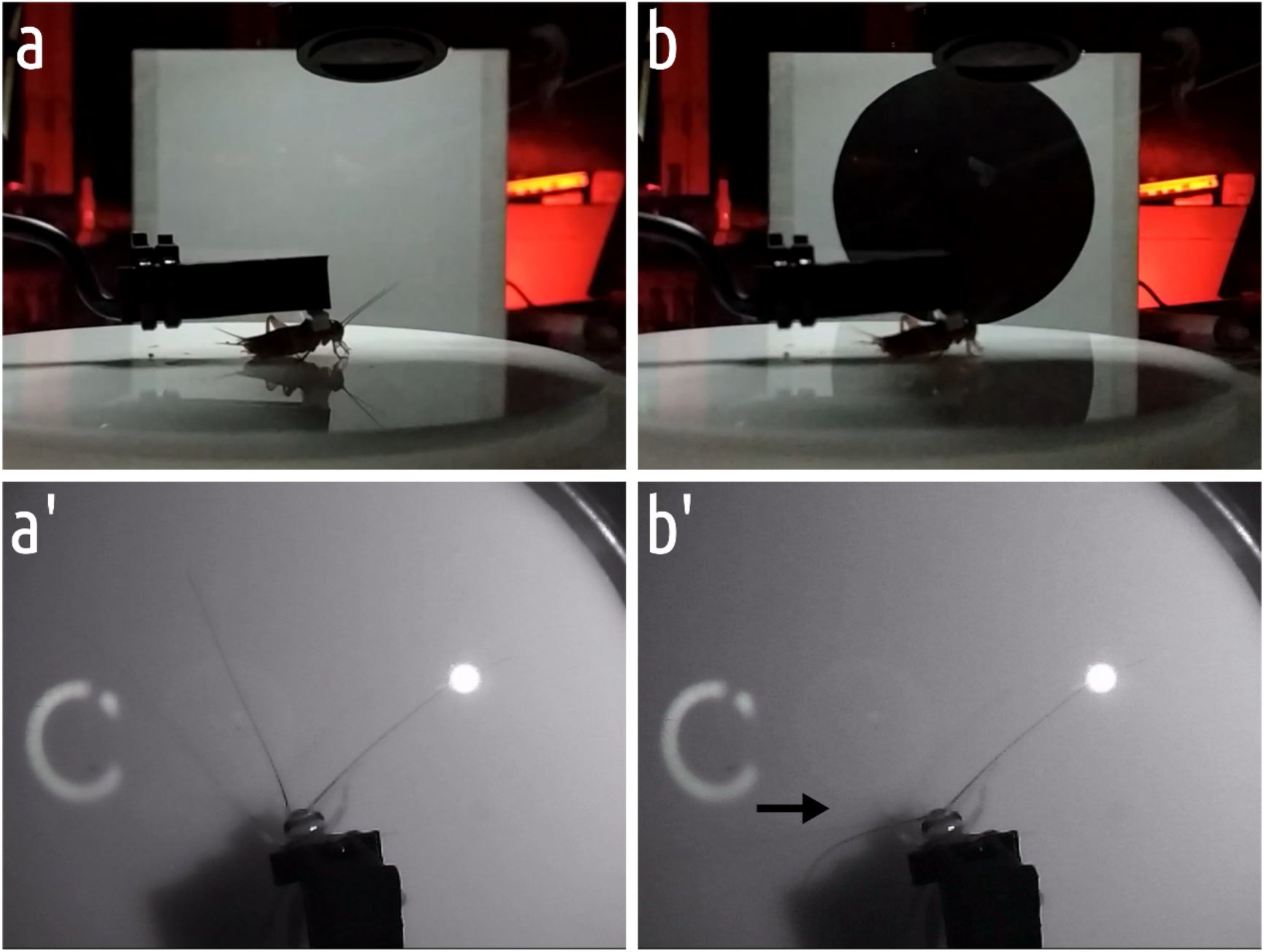
General overview of the setup (white light variant). a, b - the setup before and after the stimulus presentation. a’, b’ - the cricket’s reaction to the stimulus (direction of the stimulus indicated by the arrow).

### 2.4. Light

In order to confirm that red illumination, utilized in the main experiment, was imperceptible, and the visual cues were unavailable to insect navigating in TW setup, we utilized the well-known antennal positioning reflex towards the looming stimuli (Yamawaki and Ishibashi 2014). The tests were conducted in two variants: the first variant - under the red illumination used for the main experiment and the second - under the white light. The tethered insect was positioned on the smooth Lucite disc (acting as an omnidirectional treadmill, allowing the insect to move its legs but without providing traction sufficient for walking), and the circular looming stimulus was presented to its left side. The movement of the antennae was recorded with an infrared camera and subsequently analyzed in BehaView (v. 0.0.23; Boguszewski 2022).

### 2.5. Blinded crickets

In order to further confirm the non-visual character of observed behavior, we conducted additional control with blinded crickets. A day before the learning procedure (conducted exactly the same way as in the main experiment), the crickets were anesthetized with CO_2_, and their eyes were carefully painted over with Edding 751 paint marker (Fig. 5). This procedure did not damage the eye but rendered crickets entirely unreactive to visual stimuli.

**Figure 5.**
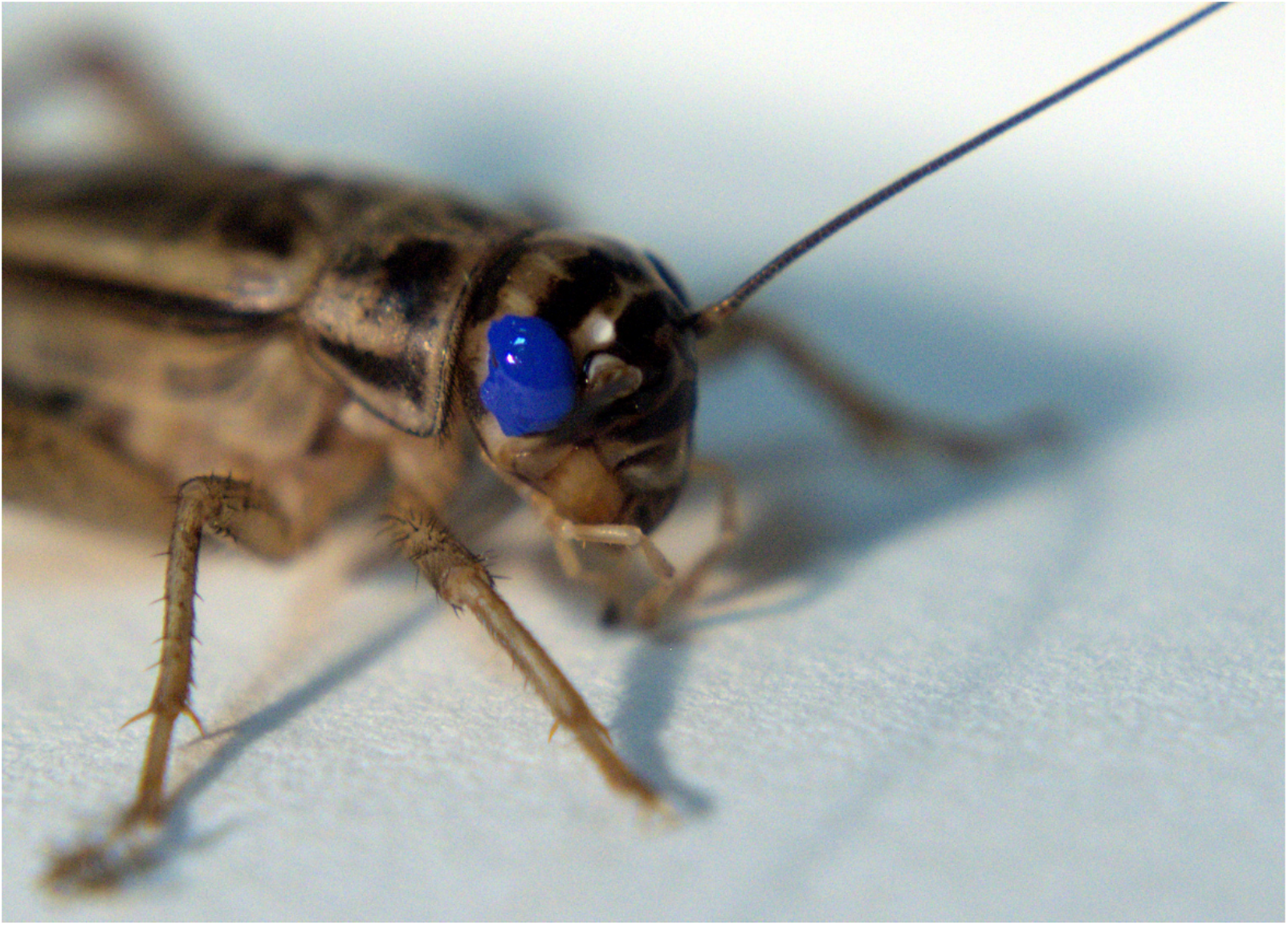
Closeup of *A. domesticus* with the painted eye. The enamel thoroughly covers the surface of the eye, impairing the cricket’s vision.

### 2.6. Center finding

Prior to the experiments, each insect was removed from the general colony and underwent an initial habituation in order to familiarize it with the transfer container (black film roll case - ø30mm, h50mm). After that, the insect was placed on the arena and left undisturbed for a 5-minute trial (Fig. 3) while a recording was captured. Between subsequent trials, the cricket was removed from the apparatus for a 5 min rest. Each session consisted of 10 trials (as in Wessnitzer et al. 2008) per arena. The procedure of each trial was analogous to a typical MWM test. The cricket was released at a random location in the arena and had to find the centrally localized, inconspicuous cool spot that allowed it to escape the aversive heat stimuli.

### 2.7. Spontaneous activity

The assessment of the spontaneous movement of crickets in all the tested arena shapes (per shape n=15, Σn=120) was performed to obtain data on how likely the crickets would be to spontaneously visit the arena center. The tests were performed in two variants - firstly, with the arena heated to the same temperature as during the main experiment, and secondly, without the heating. In both conditions, the cool spot was absent.

### 2.8. Data acquisition and processing

All recordings were captured with a Microsoft LifeCam Studio webcam and VirtualDub (v. 1.10.4.35491) software. Movement trajectories of tested insects were extracted with SwisTrack 4 software (Lochmatter et al. 2008) and further processed in R (v. 4.0.1, R Core Team 2013) with the trajr package (McLean and Skowron Volponi 2018). In order to obtain detailed data on what strategies insects used to navigate to the center, the DeepLabCut 2.0 (Mathis et al. 2018), machine learning-based tracking framework, was used. In each video, all corners (or in the case of a circular one, 4 furthermost points on the arena’s perimeter) were tracked along with insect position. This allowed the analysis of insect movement in the spatial context of the arena.

### 2.9. Statistical analysis

Statistical data analyses were performed in the R software for statistical computing (v. 4.0.1, R Core Team 2013) and GraphPad Prism 9 software. To test whether crickets showed any signs of learning to find the cool spot at arena centers, we used mixed-effects linear regression models (LMMs), utilizing the R-packages lme4 (Bates et al. 2015) and lmerTest (Kuznetsova et al. 2017). Firstly, we tested the change in the proportion of time spent on the centrally located cool spot throughout time (trial repetitions) in the different arenas. Proportion of time spent in arena center was used as response variable, which was square root transformed prior to model fitting in order to normalize the variable’s distribution. Time (trial number) and arena shape were used as predictors, also including the interaction between them, i.e., to be able to estimate different learning curves for the different arena types. Secondly, we tested the change in the latency until first arrival at the cool spot throughout time; in this model, time passed between the start of the trial and the cricket finding the cool spot was the response variable (log transformed in order to normalize its distribution), and, similarly to the previously described model, time, arena shape, and their interaction were specified as predictors. In both LMMs we used the ID of individual crickets as random effect, and also controlled for between-individual variation in slope estimates (random intercept and slope LMMs).

We used Mann-Whitney tests to compare the time proportions spent in the arenas’ central part (corresponding to occupied by cool spot) between the crickets spontaneously navigating unheated arenas and the crickets from first trial, separately for each arena type (including the tests with blinded crickets). Because of using multiple comparison tests on the same dataset, we applied Bonferroni’s post hoc P-value adjustment on the P-values from the Mann-Whitney tests, to reduce the probability of type I error.

Comparison of insects’ spontaneous activity was performed using GraphPad Prism 9 which was also used for preparation of corresponding plots. Statistical analysis consisted of two-way ANOVA, with Šidák’s multiple comparisons test

In order to investigate what strategies were utilized in order to reach the cool spot, we analyzed trajectories preceding successfully locating the cool spot (defined as the stay, lasting for at least 5 seconds spent on the cool spot). The trajectories from all trials in all arenas were analyzed, and their numerosities were counted. Each approach path, consisting of insect coordinates in every video frame during the last 3 seconds preceding the successful stay, was isolated. Paths with less than 5% time spent outside the cool spot (movement on the cool spot edge) were excluded in order to minimize noise.

For each frame the distance to the arena perimeter was evaluated to this end we employed a WKT geometry description of the and the methods from the package rgeos (Bivand and Rundel 2021). In order to compare the dynamics of approaches in all arenas, we used the distance from the arenas’ perimeters (Fig. 6). Collected wall distance time series were feature scaled and analyzed using dynamic time warping in order to extract the clusters of similar approaches. On the basis of optimization of clustering performance metrics available in the package dtwclust (Sardá-Espinosa 2019) as well as the observation of data it was decided to set the number of clusters at k=4.

**Figure 6.**
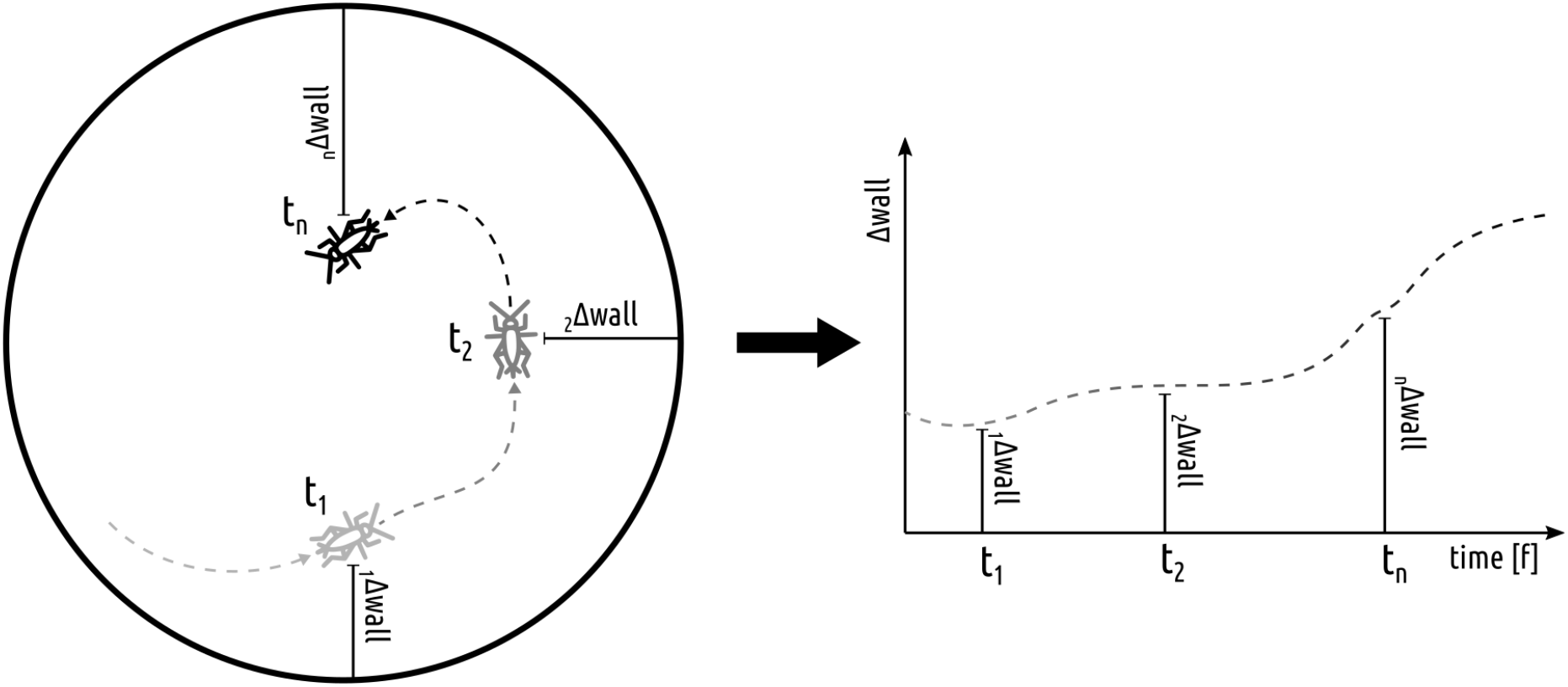
Depiction of method devised to calculate comparable one-dimensional time series from trajectories for approach patch characteristics comparison. For each timepoint (corresponding to a single video frame) of isolated approach path cricket’s minimum cartesian distance from the arena perimeter was calculated.

## 3. Results

### 3.1. Confirmation of inaccessibility of visual cues

The presence of antennal positioning reflex towards the looming stimuli starkly differed depending on under which illumination the crickets were tested (Fig. 7). Under the white illumination, the reflex was exhibited by almost all tested crickets, while under the red (used in the main experiment) illumination, it was almost universally absent, thus corroborating the inaccessibility of visual cues in the experimental setup.

**Figure 7.**
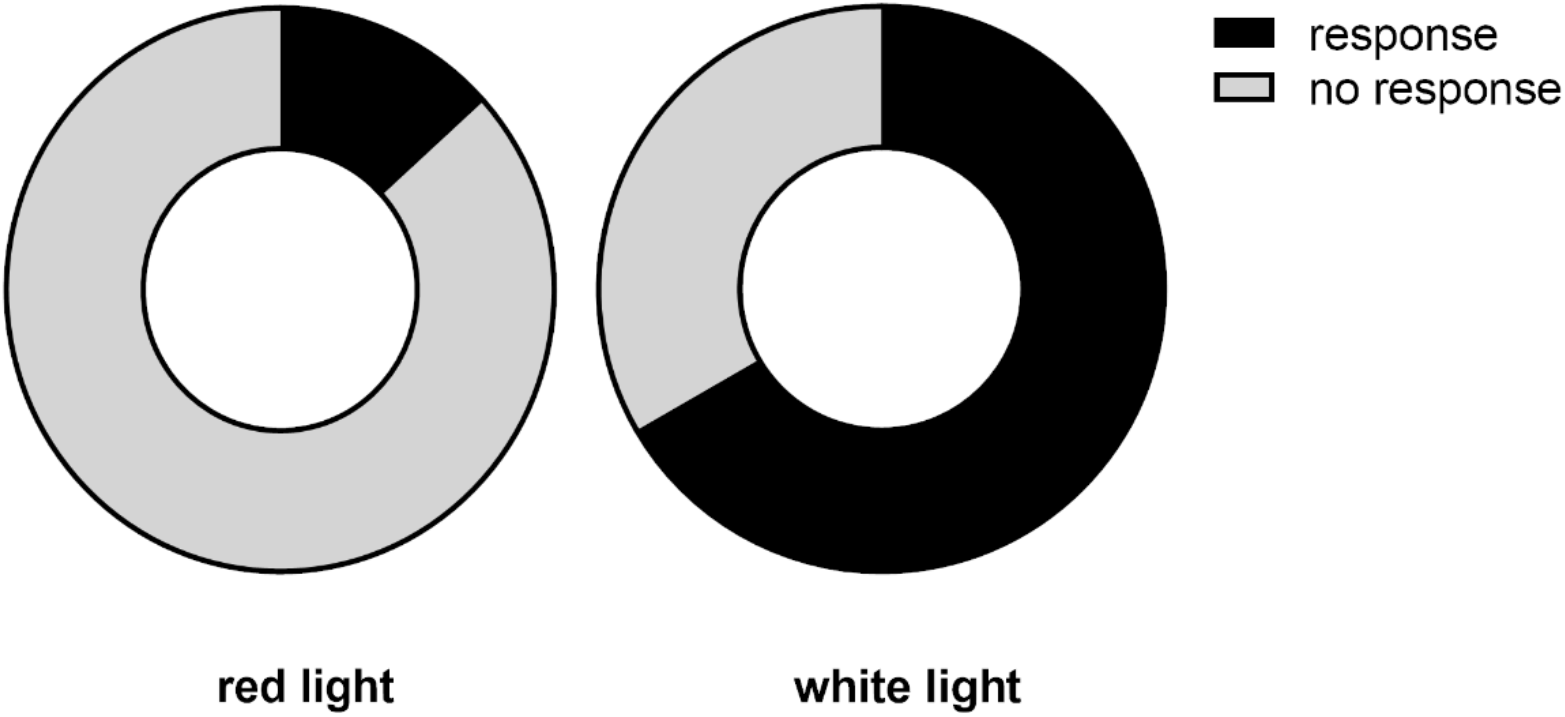
Antennal positioning response proportions to looming stimulus under two illumination variants.

### 3.2. Spontaneous activity

Patterns of spontaneous spatial exploration (Fig. 8), as indicated by the percent of time spent in proximity of the walls and spent in a centrally located spot (corresponding to the position of the cool spot in the main experiment), indicate significant differences in the exploration of various layouts. Additionally, the percentage of total time spent resting differed significantly between the circle-shaped arena and other layouts, representing the most intensive exploration of the circular arena. Considering the aims of the study, the observable differences point toward the strong thigmotaxis behavior and a strong avoidance of the central region of the arenas (especially when the surface is heated as during the main experiment), indicating the lack of initial preference of the center of any arena shape used in the main experiment.

**Figure 8.**
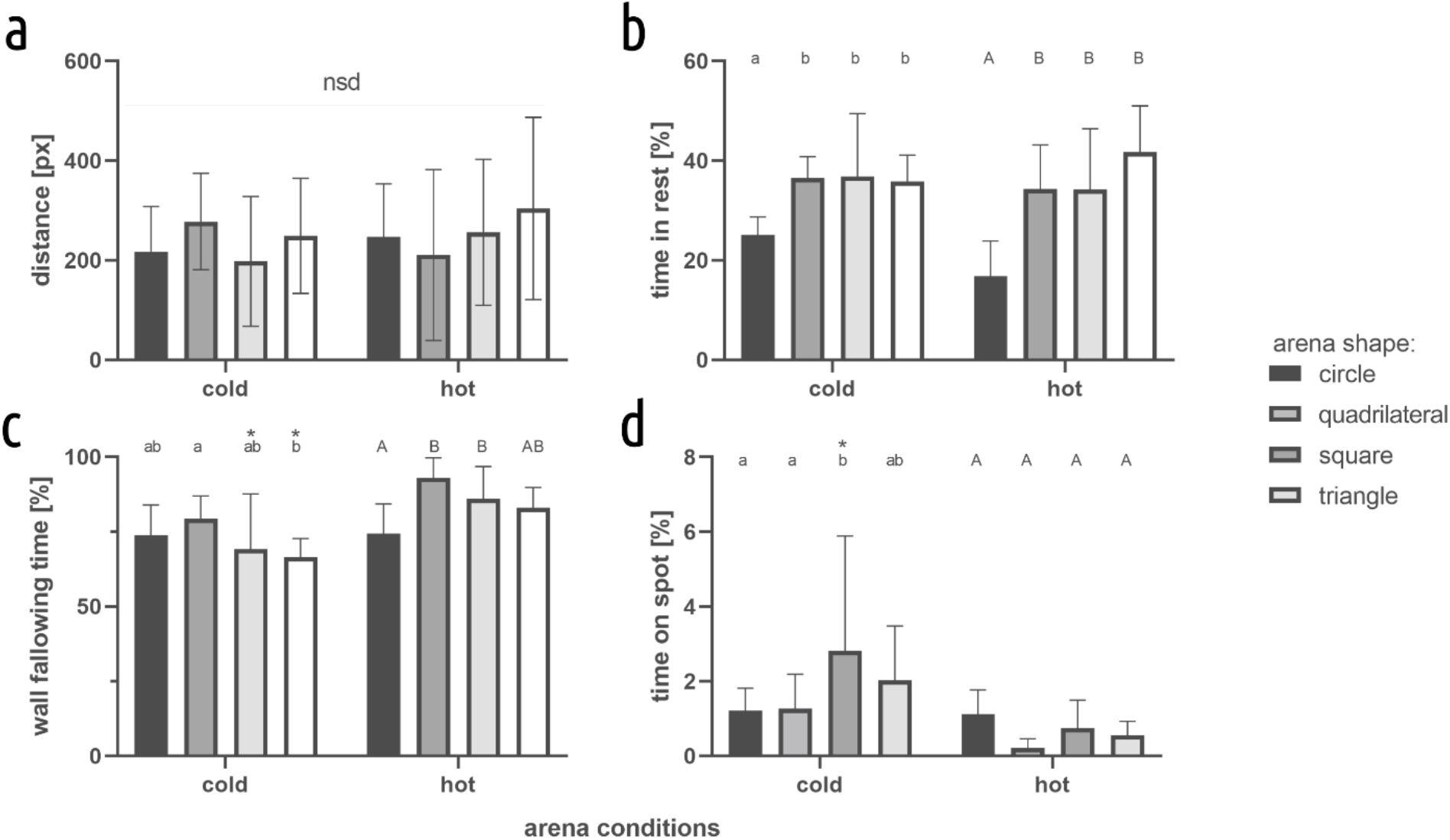
Movement characteristics for different arena shapes. a) distance, b) time in rest, c) wall following time, d) time on spot. Values are presented as mean and SD. Different letters indicate statistically significant differences between arenas - small letters for cold and big letters for hot conditions, * - statistically significant differences between conditions for particular arena shape. Two-way ANOVA Šidák’s multiple comparisons test, p < 0.05, N=15

### 3.3. Center finding learning

With the progression of trials, crickets tended to spend more time at the cool spot in all the symmetric arenas, but not in the asymmetric quadrilateral one (Fig. 9 and Table 1, Part A). We found no significant differences in the learning curves’ slopes between the circular, triangular, and square arenas. However, we found that the learning curve estimate (slope) was an order of magnitude lower in the asymmetric arena. Furthermore, with the progression of trials, crickets’ latency to find the arena centers significantly decreased in all arena types (Fig. 9 and Table 1, Part B). Arena type differences in the slopes of the latency-reduction were only found between triangular arena versus square, quadrilateral, and circular ones, as the slope of the estimated regression line was steepest in the former.

**Figure 9.**
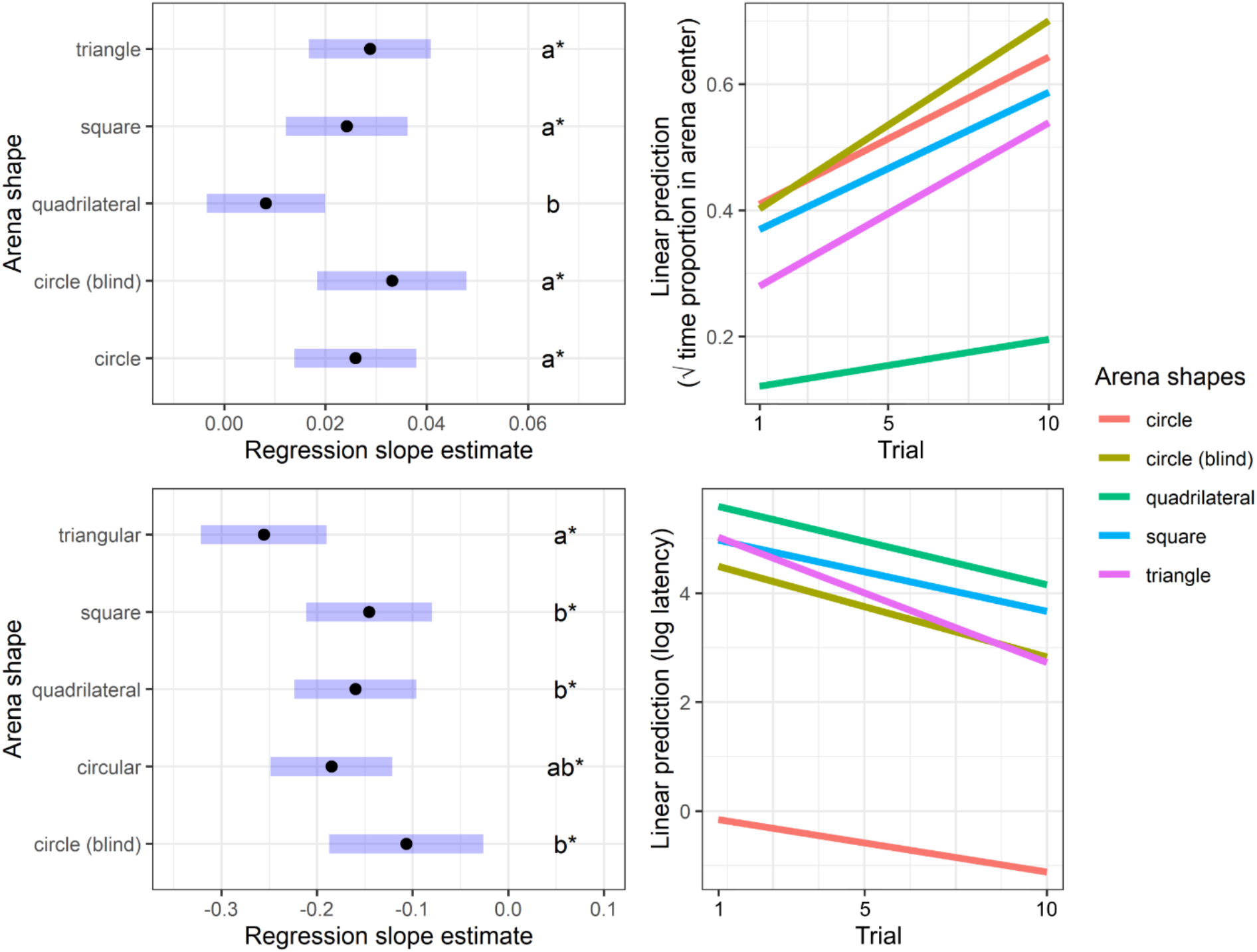
Regression slope estimates from the LMM on the time proportion spent on the cool spot (upper left) and on the latency until arriving at the arena centers (lower left) by crickets: letters denote the significance of arena type differences in slope estimates, and asterisks mark slope estimates significantly greater than zero. Regression lines are also visualized for the time proportion spent on the cool spot (upper right) and on the latency until arriving at the arena centers (lower right) as a function of time (trials).

**Table 1.**
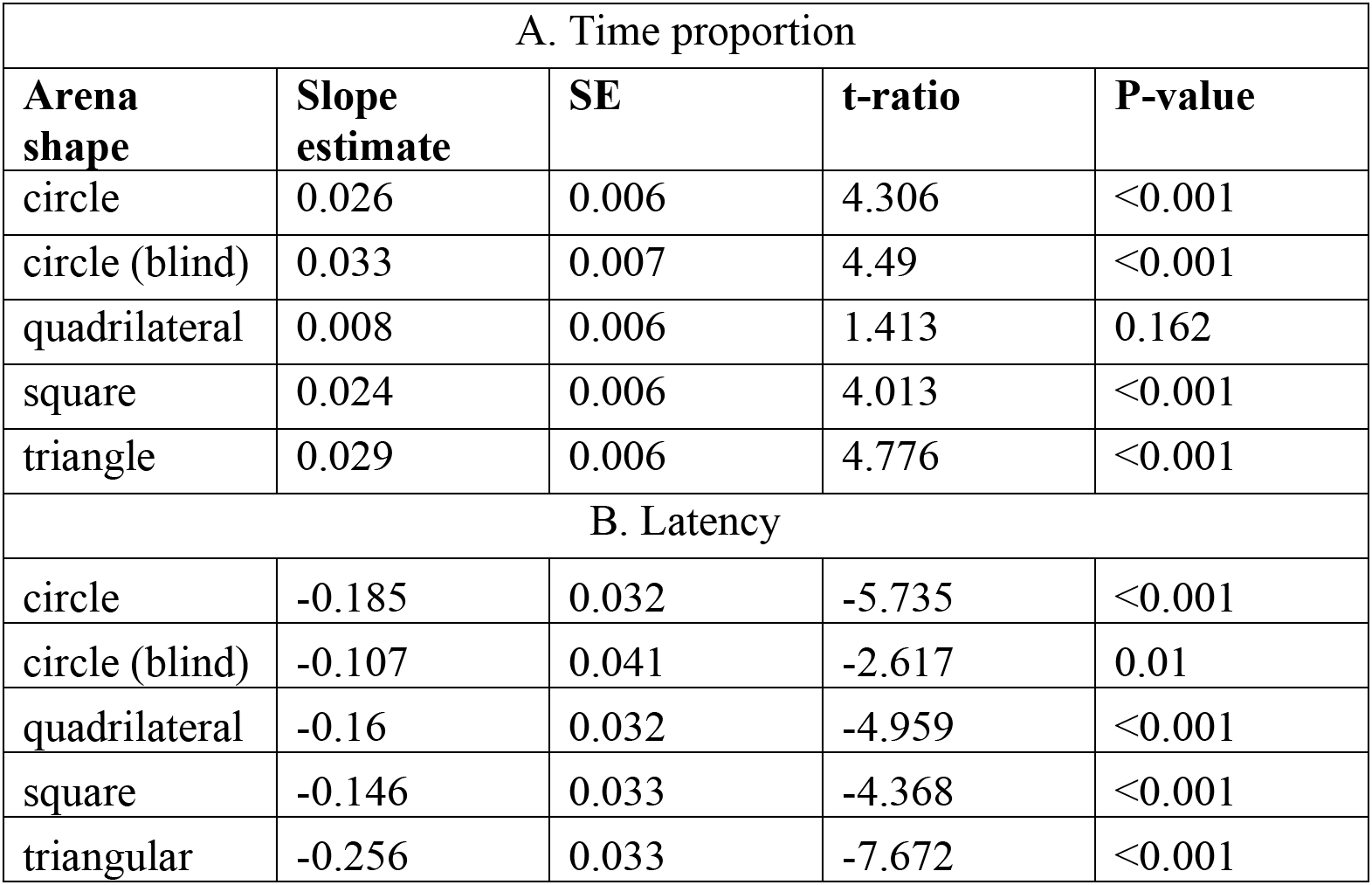
Slope parameter estimates from the LMM fitted on (Part A) the time proportion spent on the cool spot, (Part B) the latency prior to reaching the cool spot by crickets.

| A. Time proportion |  |  |  |  |
| --- | --- | --- | --- | --- |
| Arena shape | Slope estimate | SE | t-ratio | P-value |
| circle | 0.026 | 0.006 | 4.306 | <0.001 |
| circle (blind) | 0.033 | 0.007 | 4.49 | <0.001 |
| quadrilateral | 0.008 | 0.006 | 1.413 | 0.162 |
| square | 0.024 | 0.006 | 4.013 | <0.001 |
| triangle | 0.029 | 0.006 | 4.776 | <0.001 |
| B. Latency |  |  |  |  |
| circle | -0.185 | 0.032 | -5.735 | <0.001 |
| circle (blind) | -0.107 | 0.041 | -2.617 | 0.01 |
| quadrilateral | -0.16 | 0.032 | -4.959 | <0.001 |
| square | -0.146 | 0.033 | -4.368 | <0.001 |
| triangular | -0.256 | 0.033 | -7.672 | <0.001 |

### 3.4. Comparison of spontaneous exploration and performance in the first trial

Crickets spontaneously exploring unheated arenas spent significantly less time at the arena center than all groups during the first trial (Fig. 10) - circle (blind) (P < 0.001), circle (P < 0.001), triangle (P = 0.027), square, although this latter was only marginally significant after Bonferroni’s P-value adjustment (P = 0.079), with the exception of crickets navigating quadrilateral arena (P = 1).

**Figure 10.**
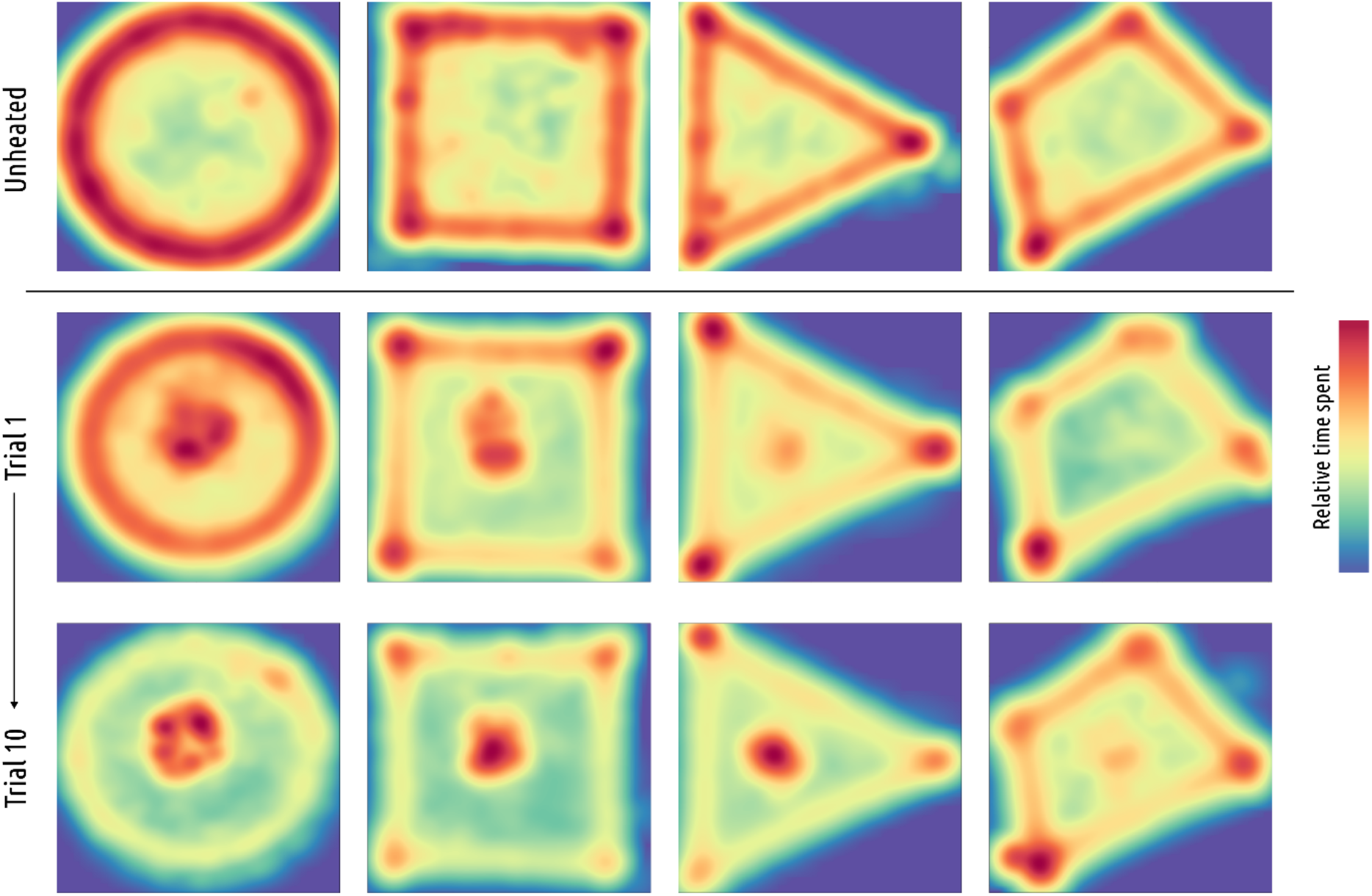
The cumulative heatmaps depict the proportion of time spent by the insects in particular places of each arena without thermal stimuli (upper row), the first trial (middle row), and the final trial (bottom row) of the study.

### 3.5. Approach paths characteristics

In total, throughout the main experiment, cricket performed 1349 successful approaches to the cool spot; out of those, based on similarity, four clusters were isolated and characterized (Fig. 11).

**Figure 11.**
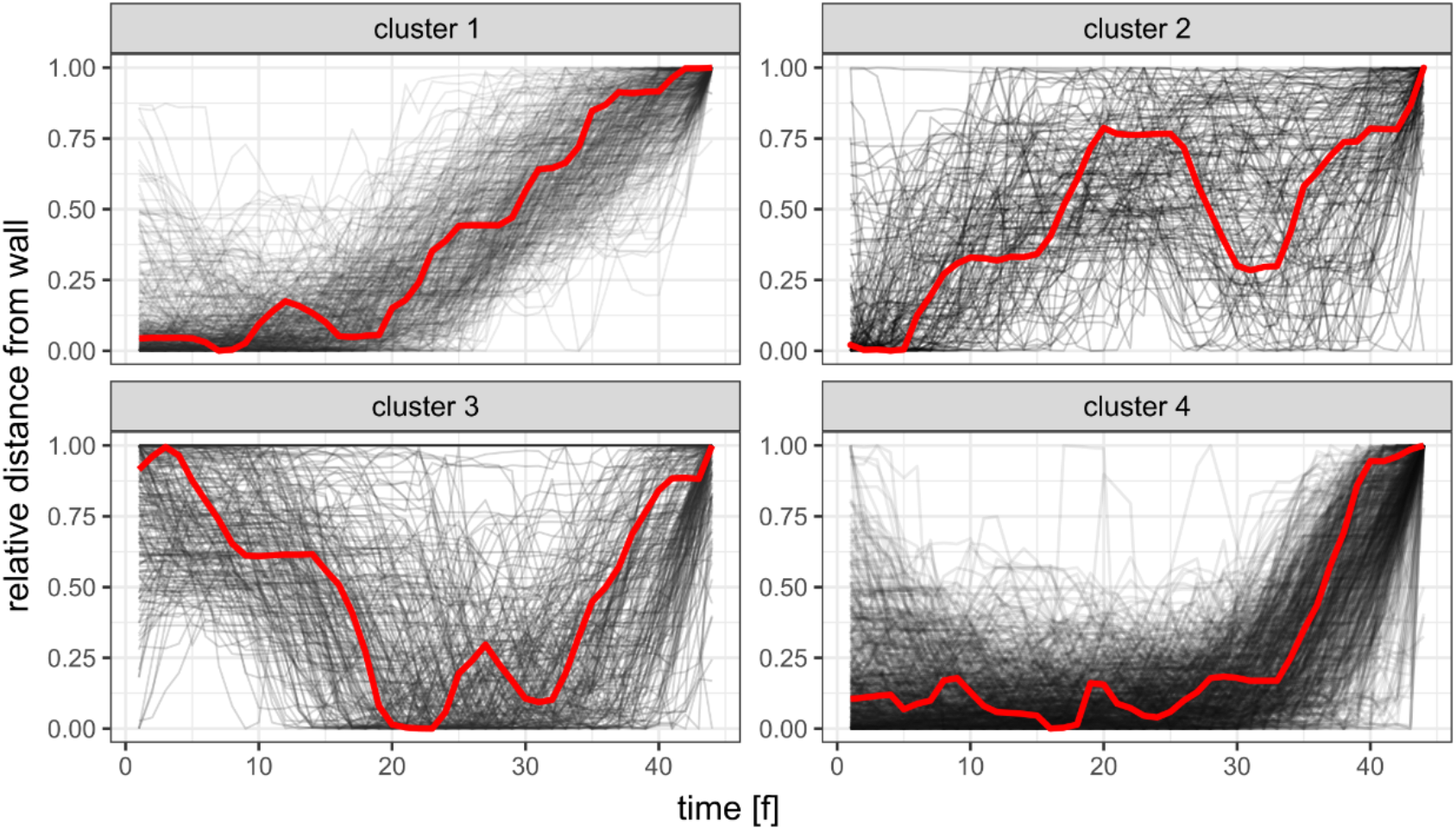
All approach paths plotted in corresponding clusters. Centroids, corresponding to most characteristic path for each cluster colored in red. Cluster members numerosities: cluster 1 - n = 326, cluster 2 - n = 151, cluster 3 - n = 228, cluster 4 - n = 608.

Approach paths numerosities in each cluster grouped by arena shapes reveal that in all arena shapes, the cluster 4 type approach paths have been used the most frequently, followed by cluster 1 type (Fig. 12). This pattern prevails even in the quadrilateral arena, with the least number of successful approaches made in total (which correspond to the observed lowest percentage of time spent on the cool spot and the highest latency to locate it). The second, less pronounced variation from the overall pattern is a slight elevation of number of cluster 3 type approaches in the circular arena.

**Figure 12.**
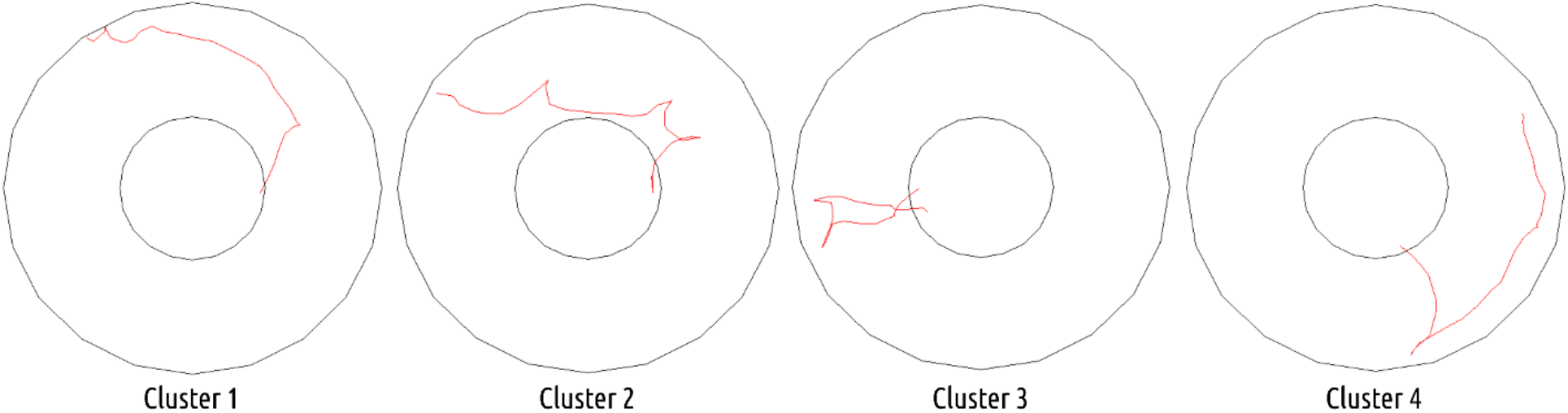
Trajectories preceding finding the centrally located cool spot from circular arena representative for each cluster. General descriptive characteristic of cluster members: cluster 1 - gradual approach towards the center both in diagonally oriented trajectories and in a spiral-like mode, cluster 2 - fast, archlike ventures towards the center and back, towards the perimeter, cluster 3 - ventures from the center, towards the walls, and back again, cluster 4 - wall-following and direct approach to center.

## 4. Discussion

In the presented study, we aimed to test whether the insects would be able to learn to successfully navigate without the visual cues, relying only on layout symmetry perceived tactilely. Additionally, we attempted to characterize the strategies allowing insects to navigate successfully, with particular attention to differentiating between heuristic ones (independent of knowledge about layout acquired by the individual) and those that relied on perception and memory of environmental layout. To this end, we employed a non-visual version of the center-finding paradigm with additional tests ensuring the inaccessibility of visual cues in experimental conditions. While we are aware that the task used in the study is exceedingly simplistic in comparison to the natural environments in which insects have to navigate in their daily life, our goal was to ensure maximal separation from the stimuli that could overshadow the possible presence of geometry-based navigation capacities in our model specie.

Despite the lack of previous insect research conducted with the center-finding paradigm, our hypotheses were driven by the general theoretical claim that sensitivity to layout geometry increases the environmental adaptation of the animals. Layout geometry generally constitutes a distinctive, robust, and computationally inexpensive cue that can be used in place finding (Cheng 1986; Gallistel 1990; Spelke et al. 2010; Thinus-Blanc et al. 2010; Tommasi et al. 2012; Hohol 2020). This claim alludes to distal evolutionary origins of sensitivity to layout geometry, which imply that it should be observed in various animal phyla. Therefore, we expected to replicate the observations previously seen in studies on vertebrates (Tommasi et al. 1997; Gray et al. 2004; Tommasi and Thinus-Blanc 2004; Tommasi and Giuliano 2014). Nevertheless, one has to remain aware of the possible variance of mechanisms among the species (Vallortigara 2018). Our hypotheses were further substantiated by previous findings that insects exhibit sensitivity to object symmetry (Giurfa et al. 1996; Møller and Sorci 1998; Rodríguez et al. 2004; White and Kemp 2020), and they can tactilely recognize previously seen objects while in the dark (Solvi et al. 2020). Furthermore, insects exhibit good navigational performance in tasks where the spatial layout is geometrically regular (Wystrach and Beugnon 2009; Sovrano et al. 2012; Lee and Vallortigara 2015). Finally, they are capable of swift conceptual learning involving the development of spatial concepts (Giurfa 2013, 2015; Avargues-Weber and Giurfa 2013).

We found that crickets can learn to localize the centrally positioned, inconspicuous cool spot. The additional tests corroborated the inaccessibility of visual cues, and thus supported the effect observed in the main experiment. The learning was significantly more efficient in all the symmetric arenas (circular, square, and triangular) in comparison to the asymmetric quadrilateral one. More specifically, the learning curve estimates were significantly higher in the symmetric arenas than in the asymmetric one, and the latency of finding the center was significantly longer in the latter. This effect was indicated by a higher intercept estimate in the regression model. Nevertheless, in subsequent trials (in all the arenas), the time spent in the center increased and the latency decreased, albeit, in the asymmetric arena, the effect was less pronounced. Furthermore, in all the symmetric arenas, the number of successful approaches was substantially higher in comparison to the asymmetric quadrilateral one. The data obtained from the spontaneous exploration conditions emphasizes the learning aspect of the observed behavior as the cricket, aside from learning to find the center, had to suppress its thigmotaxis reflex. The spontaneous visits to the center of the arenas were rare in both (heated and unheated) conditions. Aversion to open spaces exhibited by crickets also could account for crickets leaving the cool spot after discovering it. While the crickets’ mode of spontaneous exploration of the arenas differed, those differences do not seem to translate to the results of the learning trials. Furthermore, while the presence of the corners significantly increased the time spent by the crickets in the proximity of the walls, it was not significantly different in the quadrilateral arena. This data supports the observation that differences in learning could not be explained only by the preference for sharp corners. Additionally, overall exploratory activity (as indicated by the traveled distance) does not seem to be related to the complexity of the arenas. Overall, the results corroborate our hypotheses, allowing us to infer that layout symmetry facilitates spatial learning and can be considered a viable cue for successful place finding in insects.

A significant reduction in observed latency indicates that the cricket’s capacity to find the center could not be explained entirely in terms of learning to interrupt the random search (or scanning) when the cool spot is reached. If this were so, the time spent on the cool spot would increase, but the latency would have remained constant (see Foucaud et al. 2010). Furthermore, another study performed using the MWM test revealed that executing heuristic search routines, such as scanning or chaining, could lead to comparable latencies as direct search and related strategies (Wolfer and Lipp 2000; Garthe et al. 2009). Our approach path analysis revealed that, indeed, both heuristic and directed strategies are employed (Fig. 11, Fig. 12), however, with substantial dominance of a semi-directed (thigmotactic phase preceded the direct navigation to the target spot) in all the arenas (Fig. 13). Before most of the successful navigation bouts insects tended to visit the perimeter of the arena and subsequently travel directly to the cool spot (Fig. 12). We consider that aforementioned thigmotactic phase preceding the travel to cool spot could be interpreted as an orientation period used to adjust the memorized allocentric layout model. As the employed experimental paradigm is devoid of visual cues, insects had to rely solely on tactile cues, in contrast to, e.g., rats in MWM, which could instantaneously access visual information about their position.

**Figure 13.**
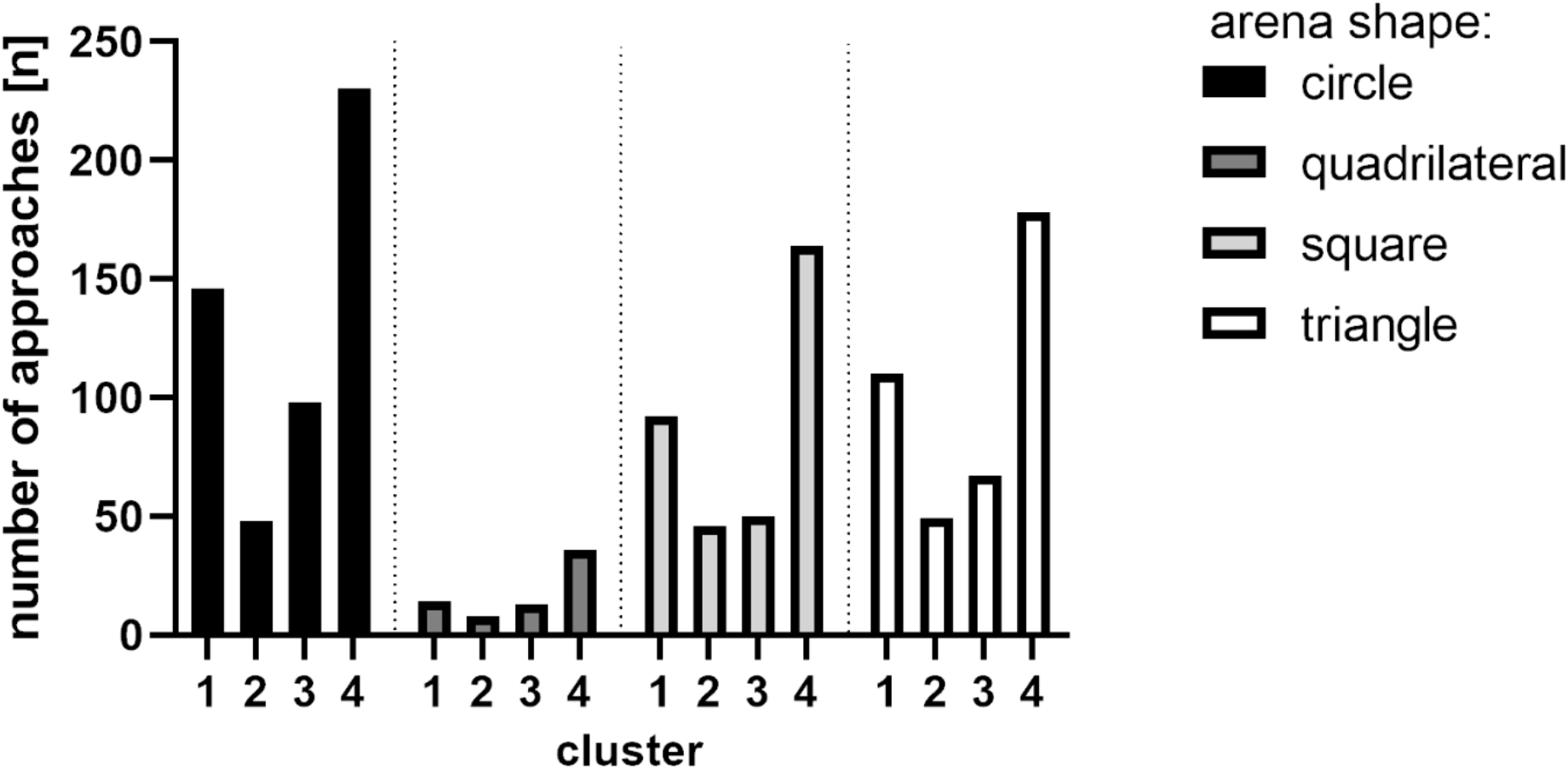
Total numerosity of approach trajectories corresponding to each cluster in all arenas.

By its distal nature, the visual information provides an overview of a much larger area. This result may be considered supporting evidence for the memorizing arenas’ layouts by insects (Gould 1986; Wehner and Menzel 1990; Webb 2019). Furthermore, as was shown in the 3rd cluster, insects were able to return to the cool spot without performing the extensive thigmotactic orientation phase. Therefore, it seems that aside from allocentric memory of arena layout, insects were able to retain egocentric memory of cool spot location and utilize it to navigate back in case of brief detours from the cool spot. Approach paths grouped in clusters 1 and 2 highly resemble non-directed, heuristic search strategies, namely chaining and scanning. They consist of repeatable movement patterns, respectively spiral-like and consisting of arch-like detours from the wall, which are executed until the cool spot is reached. Interestingly, those strategies were rarely utilized in the asymmetric arena, and out of the few animals that managed to locate the cool spot, most were using semi-directed strategy.

This indicates that, in principle, it is possible for crickets to successfully navigate in asymmetrical arenas, though for some reason, it is exceedingly more challenging. We speculate that the informational complexity of the arenas may explain one probable cause of this effect. Symmetric arenas are computationally easier to encode (e.g., a circular arena could be described only by its radius, all the walls and angles of a square and triangle arenas are equal) in contrast to the asymmetric quadrilateral arena (in which all walls lengths and angles differ). On the contrary, crickets in the wild have to navigate in much more complex environments than simple testing arenas. However, in natural environments, they are constantly provided with information from more than one modality, and some results suggest that multimodal information may facilitate spatial learning (Taevs et al. 2010; Hebets et al. 2014; Buehlmann et al. 2020; Sun et al. 2021). Such possible information processing could be executed by the central complex as was proposed by Xuelong Sun et al.’s (2021) model. The central complex receives projections from antennal lobes, which are known to process, aside from chemosensory information, also the tactile stimuli (Nishino et al. 2005). Hence, we propose that in the face of such limited cues, encoding the asymmetric quadrilateral arena could exceed the working memory capacity of most of the crickets, thus preventing them from successfully orienting in the arena during the thigmotactic phase.

### Concluding points

- Crickets are capable of learning to localize the centrally positioned, inconspicuous cool spot.
- The symmetry of the arena significantly facilitates crickets’ learning to find a cool spot.
- Crickets used both heuristic and directed strategies of approaching the target, with the dominance of a semi-directed strategy (thigmotactic phase preceding direct navigation to the target).
- We hypothesize that the poor performance of crickets in the asymmetrical quadrilateral arena may be explained by the difficulty of encoding its layout with cues from a single modality.
- We propose that further exploration of observed effects may be followed by testing the spatial learning in other arenas, rectangular or rhomboidal ones.
- The possible involvement of informational inputs from antennal lobes in navigation in the presented task could be studied with experiments involving either removal of the antennae or lesions to antennal lobes.

## Acknowledgements

This study was funded by the Polish Ministry of Science and Higher Education (the Diamond Grant program, id: DI2015025445, PI: Bartosz Baran). Authors would like to thank Barbara Webb and Antoine Wystrach for their comments on experimental design and preliminary results, Véronique Izard and Nora Newcombe for helpful remarks regarding the analyses and conceptual framework, and Rafał Czajkowski and Jan Manuel Rodriguez Parkitna for reading and commenting on the very early drafts. Moreover, we would like to thank Ken Cheng and the three anonymous Reviewers who provided insightful comments on the previous version of this manuscript and encouraged us to include auxiliary control conditions as well as the approach trajectories analysis. We also thank Sylwia Butkiewicz for reading the final draft.

## Contributions

Bartosz Baran and Mateusz Hohol wrote the manuscript. All authors commented on the manuscript. Bartosz Baran, Michał Krzyżowski, Jacek Francikowski, and Mateusz Hohol conceived, designed, and performed the experiment. Bartosz Baran, Zoltán Rádai, and Jacek Francikowski analyzed the data and prepared the figures and tables. All authors substantially contributed to the interpretation of the data. All authors read and approved the submitted manuscript.

### Ethical approval

All applicable international, national, and institutional guidelines for the care and use of animals were followed. The methods and procedures conformed to guidelines of the Faculty of Natural Sciences of the University of Silesia in Katowice for testing invertebrate animals.

### Data availability

Raw data are available at the Open Science Framework: https://osf.io/w7db2/. All the further information necessary to replicate the study and its results is available from the corresponding authors upon request.

